# Opposing Regulation of Tau Condensation by Phosphorylation Levels and Molecular Chaperone Interactions

**DOI:** 10.64898/2026.10.27.762205

**Authors:** Cecilia Mörman, Tanushree Agarwal, Lei Ortigosa-Pascual, Tuomas PJ Knowles, Axel Abelein

## Abstract

Aggregation of the microtubule-stabilizing protein Tau into insoluble deposits is characteristic for several neurodegenerative diseases, including Alzheimer’s disease. While phosphorylated Tau (pTau) appears as a normal post-translational modification, the accumulation of hyperphosphorylated Tau aggregates represents a hallmark of these disorders. The molecular mechanisms involved in the conversion from functional Tau into pathological assemblies are still not elucidated. Here, we identified the role of phosphorylation levels on the phase separation and aggregation behavior of Tau, and how these processes can be counteracted by the molecular chaperone-like Bri2 BRICHOS domain. We found that the phosphorylation pattern modulates Tau fibrillation and enhances its droplet-formation propensity. Remarkably, Bri2 BRICHOS generally shifts Tau phase boundaries, counteracting the dense state, and inhibits Tau fibril formation, irrespectively of the degree of phosphorylation. Solution NMR results pinpointed the binding site of Bri2 BRICHOS on pTau, both in the dilute phase and droplet state, revealing an effect on specific regions and phosphorylated residues, which appear to be crucial for the formation and maintenance of the droplet state. Taken together, this study uncovered the opposing regulatory roles of phosphorylation and the anti-amyloid chaperone Bri2 BRICHOS on biomolecular condensation and fibrillation of Tau, providing detailed insights into the molecular mechanisms involved in tauopathies, which may facilitate future treatment strategies using chaperone-based inhibitors.

## INTRODUCTION

Reversible post-translational modifications (PTMs) are important regulatory processes in cells. In fact, it has been estimated that 70% of the proteins in eukaryotic cells are regulated by phosphorylation^1^ with a kinome of 518 different kinases^2,3^. Abnormal phosphorylation reactions may however contribute to severe disease developments. Neurodegenerative diseases, such as Alzheimer’s disease (AD) and tauopathies, are irreversible conditions affecting the nervous system and were histologically characterized by Tau protein deposits of amyloid material^4^. PTMs affect the function, aggregation, and degradation of Tau, and common PTMs include phosphorylation, acetylation, methylation, among others^5^. The molecular details of how functional Tau protein transitions into the aberrantly aggregated and phosphorylated states found in disease remain unknown.

Tau is an intrinsically disordered protein with a physiological function as a microtubule-stabilizing protein, as well as interactions with other biomolecules such as RNA^6,7^. Tau can phase separate into dynamic liquid droplets, also called biomolecular condensates, and such droplets have emerged as an important part of Tau physiology with potential connection to pathology where early assemblies of Tau within droplets may serve as precursors of aggregation into amyloid fibrils^8–10^. Native Tau forms droplets at physiological concentrations, and the critical concentration for phase separation is modulated by the degree of phosphorylated residues^11,12^. Phosphorylation of Tau decreases its affinity for microtubules and is considered a normal regulatory process under healthy conditions^13,14^. In contrast, hyperphosphorylated Tau deposits are found in disease conditions, and whether the abnormal protein behavior originates from abnormal hyperphosphorylation, from phosphorylation of already-aggregated protein, or a combination of the two processes, remains debated.

The definition of hyperphosphorylation varies in the literature, ranging from 20 to 44 phosphorylated residues^11,13,15^, or expressed as phosphate-to-protein ratio of 2-3 for healthy versus 6-8 for pathological conditions^11^. Tau has 85 putative phosphorylation sites, of which about 50 have been found to be modified^13^. The full-length Tau protein (2N4R isoform with 441 residues) has a positive net charge at physiological pH with negatively charged N- and C-termini flanking a positively charged middle segment that covers the P1, P2, and microtubule-binding (MTB) repeat-region domains (Fig. 1A). Each phosphorylated residue adds a charge of −1 to the overall protein net charge, considerably changing the protein’s properties.

**Figure 1.**
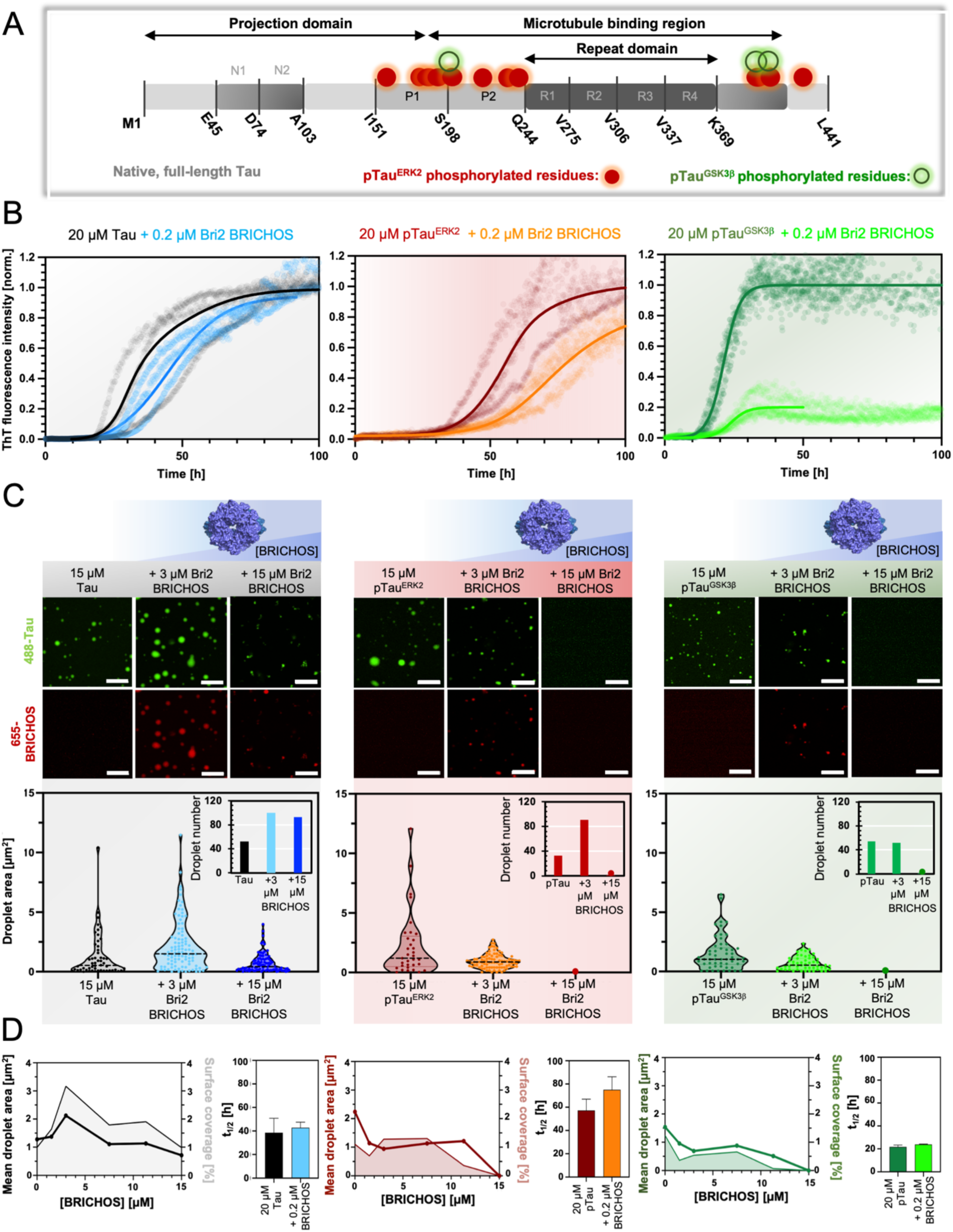
Bri2 BRICHOS inhibits pTau droplet formation and fibrillation. **(A)** Schematic presentation of native Tau and pTau, with phosphorylated residues highlighted in green for GSK3β and red for ERK2 kinases. **(B)** Aggregation kinetics of 20 μM Tau and pTau in the absence or presence of 0.2 μM oligomeric Bri2 BRICHOS in 20 mM NaP pH 7.4 buffer applying continuous agitation. The kinetic curves are normalized to the condition without Bri2 BRICHOS. **(C)** Fluorescence microscopy images of 15 μM Tau or pTau with 3 or 15 μM oligomeric Bri2 BRICHOS in droplet-promoting buffer of 8% PEG8000 and 20 mM NaP pH 7.4. 1% Atto-655 Bri2 BRICHOS and/or 1% Atto-488 Tau were supplemented to the samples to visualize the presence of droplets. The complete concentration series is presented in SI Figure S2. The scale bars represent 10 μm. **(D)** Quantitative analysis of droplet properties shown in (B), (C), and SI Figure S2, comparing the mean droplet area (bold and circles) and total surface coverage of droplets (line and colored area) as a function of Bri2 BRICHOS concentration (left panel for each protein), and aggregation halftimes where the error bars refer to the standard deviation of four technical replicates (right panel).

Multivalent and electrostatic interactions are key for the phase separation of Tau, and changes of the net charge towards neutral could potentially decrease the droplet propensity and instead increase the aggregation propensity due to an attenuation of intra- and/or intermolecular electrostatic repulsion^16,17^. An extended Tau conformation has been suggested for native Tau condensates^18^. A recent study reported increased flexibility and extended conformation for monomeric phosphorylated Tau (pTau), owing to the introduced electrostatic interactions from the phosphate groups, which results in increased intermolecular interactions for phase separation^11^. Moreover, potential biomarkers utilize certain variants of phosphorylated Tau, such as pTau217, pTau231 and pTau181, in body fluids of cerebrospinal fluid and blood, serving as a diagnostic and potentially prognostic tools^19^. The pattern of phosphorylated sites thus appears to be important for disease initiation and/or progression as well as for diagnostic biomarkers.

The typical Tau fibril core is surrounded by a “fuzzy coat” consisting of largely unstructured N-and C-termini^20^, within many of the phosphorylated residues are located. The fibril core itself is not sufficient to replicate AD-like seeding behaviour associated with disease mechanisms^21^, suggesting the importance of the unstructured parts and PTMs for disease progression. Notably, fetal Tau, a shorter variant of the full-length Tau protein and one of the six isoforms present in the human brain, has a phosphorylation pattern resembling that reported for AD brains^13,22^. Fetal Tau is, however, not toxic for the developing brain, and it has been suggested that this pattern may be important for growth or regeneration of the brain cells^23–25^. Why this phosphorylation pattern is essential during development yet associated with pathology during ageing, and how such adverse outcomes might be counteracted, remain open questions that merit further investigation.

Several different strategies have been applied to combat protein aggregation in diseases^26,27^, ranging from passive immunization to stabilization of proteins involved in proteolytic cleavages or kinase/phosphatase activity, to disaggregation of already-formed aggregates^28^. A recently suggested promising approach relies on molecular chaperones or chaperone-like proteins to prevent aggregated and potentially toxic species^29–31^. Bri2 (integral membrane protein 2B) is a type II transmembrane glycoprotein expressed in various tissues, including the brain^32^. Like Tau, phosphorylation modifications of Bri2 lead to functional changes, in this case, an increase in neuronal outgrowth and differentiation^33^. In AD, altered Bri2 concentration and activity have been reported in the hippocampus^34^. Bri2 exhibits hydrophobic and aggregation-prone regions and carries a BRICHOS domain that acts as an internal chaperone for the pro-protein, preventing misfolding of the aggregation-prone parts. The BRICHOS domain is approximately 100 residues in size, is found in at least 10 protein families in humans^32^, and BRICHOS from the Bri2 protein passes the blood-brain barrier and exhibits anti-amyloid activity both *in vitro* and *in vivo* against several proteins involved in neurodegenerative disease, including amyloid-β, Tau, α-synuclein, and IAPP^35–43^. Remarkably, Bri2 BRICHOS also modulates the phase separation properties of native Tau^41^. Bri2 BRICHOS adopts distinct functional assembly states, ranging from monomer to high-molecular-weight oligomers and, unlike native Tau, is negatively charged^35^.

Mechanistic details into Tau protein dysfunction, including disease-related phosphorylation, are of particular importance for preventing Tau-related neurodegeneration and for designing therapeutics that selectively target disease-related Tau states^44^. Here, we identified how the phosphorylation patterns of different pTau variants determine the phase space and aggregation kinetics and how these processes can be opposingly regulated by the chaperone-like Bri2 BRICHOS domain.

## RESULTS

### Bri2 BRICHOS inhibits pTau protein fibrillation

Two differently phosphorylated Tau protein variants were obtained by phosphorylation reactions *in vitro* using two endogenous serine-threonine kinases; mitogen-activated protein kinase (ERK2) and glycogen synthase kinase 3β (GSK3β). Both enzymes show ubiquitous expression in the brain and have previously been described as important regulators of Tau phosphorylation, associated with neurodegeneration^23,45,46^. *In vitro* phosphorylation of Tau with ERK2 gave rise to a major population with at least 10 confirmed phosphorylated residues, whereas GSK3β phosphorylation resulted in at least 3 phosphorylated residues (Fig. 1A and SI Fig. S1). The two pTau variants in combination with non-phosphorylated native Tau gave a system with a positively charged Tau (native), a variant with a net charge around zero (pTau^GSK3β^), and another one with a negative net charge (pTau^ERK2^) at physiological pH 7.4.

To gain insights into the pTau protein aggregation behavior, we conducted aggregation kinetics assays using 20 μM pTau and native Tau proteins. The assays were performed without co-factors in the buffer system using agitation with small glass beads^41,47^. Alike native Tau, the pTau variants formed ThT-active aggregates over time, where pTau^GSK3β^ aggregated faster (aggregation halftime t_1/2_ ∼ 21 h) than native Tau (t_1/2_ ∼ 38 h) and pTau^ERK2^ (t_1/2_ ∼ 57 h) (Fig. 1B, D right panels), in line with similar aggregation behaviour reported in the literature^23^. To investigate the effect of Bri2 BRICHOS we used the Bri2 BRICHOS R221E mutant^38,41,48^, since this variant was previously applied in *in vivo* studies^42,43^ and the inhibitory effect was characterized in great detail^38,41,48^. Further, the oligomeric fraction of Bri2 BRICHOS R221E (hereafter referred to as Bri2 BRICHOS) has previously been shown to be superior in inhibiting native Tau aggregation and hence this assembly state was predominantly used in this study. When conducting aggregation kinetics assays in the presence of 0.2 μM Bri2 BRICHOS, we observed that the fibrillation was delayed for all protein variants. Collectively, Tau forms ThT-active aggregates and Bri2 BRICHOS can inhibit fibrillation irrespectively of the phosphorylation pattern.

### Bri2 BRICHOS shifts mesoscopic phase separation of pTau variants

To further examine the Bri2 BRICHOS modulating effects of phosphorylated Tau proteins and their phase separation behavior, we applied fluorescence microscopy and initiated droplet formation by using 8% PEG8000. 15 μM of pTau^ERK2^, pTau^GSK3β^, and native Tau supplemented with 1% Atto-488-fluorescence label readily formed molecular condensates at time zero (Fig. 1C and SI Fig. S2). We observed that the droplets fused and grew in size upon maturation (SI Fig. S3), indicative of liquid-like condensate behavior. The droplet propensity for all proteins was overall in a similar range, showing the highest droplet propensity for pTau^ERK2^, followed by pTau^GSK3β^ and native Tau (Fig. 1C and D left panel). Since pTau^ERK2^ exhibits more phosphorylation sites, and hence a strong modulation of the net charge upon phosphorylation, the Tau droplet propensity is seemingly dependent on the net charge.

Addition of micromolar concentrations of Atto-655-Bri2 BRICHOS resulted in Bri2 BRICHOS incorporation into the droplets (Fig. 1C), in line with earlier studies for native Tau^41^. Adding Bri2 BRICHOS to a concentration of 3 μM led to an increase in droplet size, number, and total coverage surface for native Tau (Fig. 1C, D left panel). Further addition to equimolar concentrations results in that the droplet formation propensity was reduced but droplets were still present (Fig. 1C, D left panel). For pTau, an inhibition of pTau droplet formation was evident upon addition of Bri2 BRICHOS, with a reduction of the droplet size apparent for conditions at 1:5 Bri2 BRICHOS:pTau ratio toward a complete dissolution of pTau condensates at equimolar concentrations (Fig. 1C, D left panel). Before dissolution, distorted droplets or percolated clusters were observed. Remarkably, the Bri2 BRICHOS inhibitory effect was more prominent for both pTau variants compared to native Tau (Fig. 1D left panel). After maturation for 24 h, smaller droplets were observed in the presence of low Bri2 BRICHOS concentrations, while the percolated clusters at higher Bri2 BRICHOS concentrations remained (SI Fig. S3). Qualitatively, dilution of the mature droplet samples resulted in dissolution of the droplets, resulting in fewer and smaller droplets, which is indicative for reversible and still liquid-like droplets (SI Fig. S4).

Given that condensate and fibril formation may be either dependent or independent processes, governed by the conditions^49,50^, here we found that pTau^GSK3β^ more readily formed amyloid fibrils, yet with lower droplet formation propensity compared to pTau^ERK2^ (Fig. 1). Remarkably, Bri2 BRICHOS generally inhibits both fibril and droplet formation for these tested phosphorylated Tau variants, irrespectively of the specific phosphorylation pattern.

### pTau phase separation is modulated by the phosphorylation pattern and Bri2 BRICHOS

To further explore Bri2 BRICHOS’ effects on pTau phase separation throughout the phase space, we generated high-resolution phase diagrams for pTau^GSK3β^, pTau^ERK2^, and native Tau across a range of different PEG8000 concentrations, both in the absence and presence of Bri2 BRICHOS, using the PhaseScan platform^51,52^ (Fig. 2). PhaseScan is a droplet-based microfluidic approach that produces thousands of droplets, enabling rapid and quantitative mapping of phase behaviour across multidimensional chemical space.

**Figure 2.**
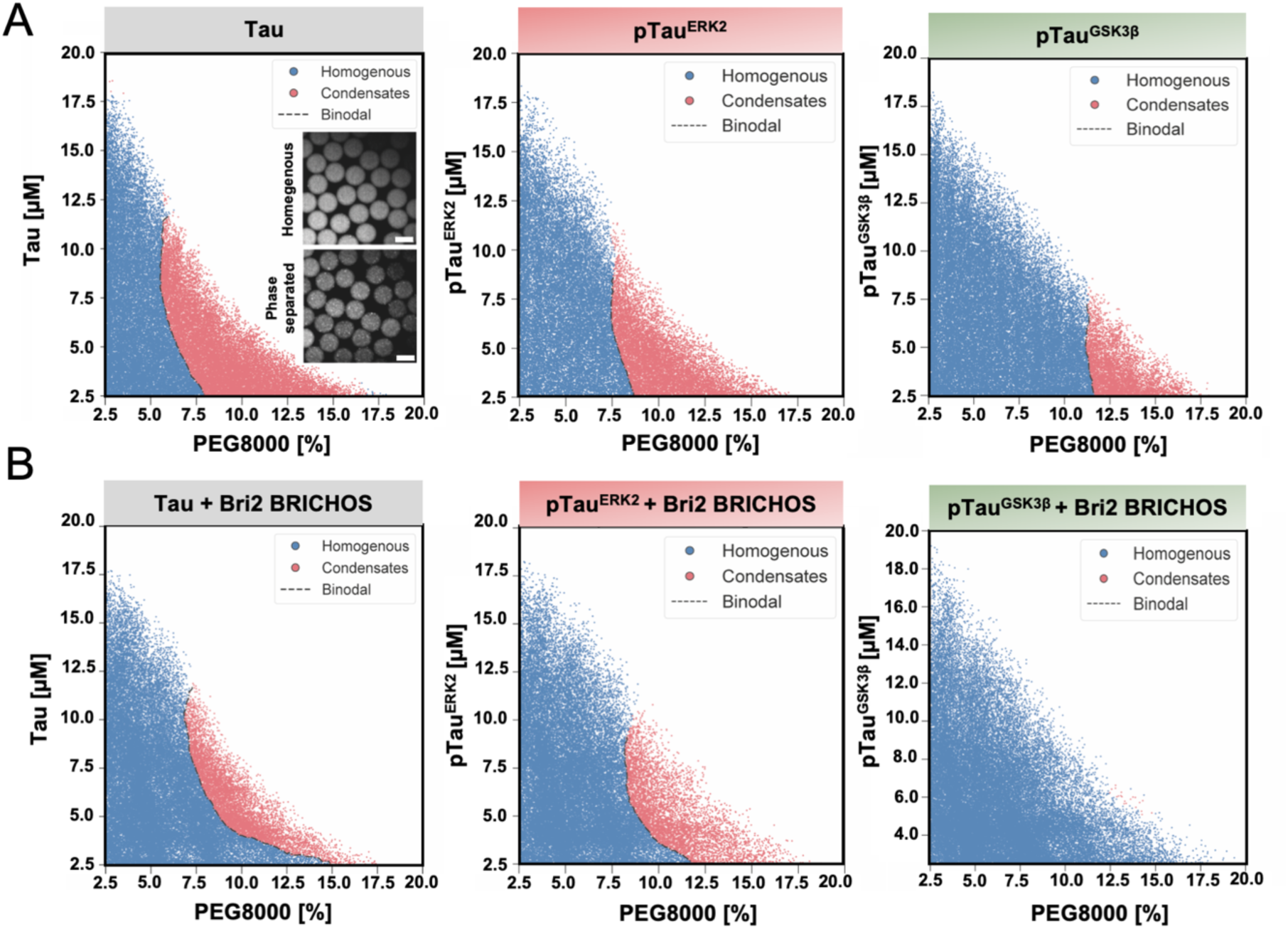
Modulation of phase diagrams of Tau and pTau by Bri2 BRICHOS. High-resolution phase diagrams generated using PhaseScan for Tau, pTau^ERK2^, and pTau^GSK3β^ as a function of PEG8000 concentration **(A)**, and the effect oligomeric Bri2 BRICHOS **(B)**. Each point represents an individual microdroplet, classified as homogeneous (blue) or phase-separated (red). Dashed lines indicate the inferred binodal phase boundaries. The number of analyzed microdroplets per condition in (A) from left to right is n = 44994, n = 36522, and n = 50055, and in (B) n = 49398, n = 44052, and n = 51541. Representative widefield images are displayed; scale bar 100 µm.

Briefly, aqueous solutions containing fluorescently labeled Tau (Alexa Fluor 488), Bri2 BRICHOS (Atto-Flour 647), and PEG8000 (Alexa Fluor 546) were combined within a microfluidic device to generate droplets spanning a continuous range of component concentrations. Each component was uniquely barcoded by fluorescence, allowing absolute concentrations within individual droplets to be determined via calibration against known standards. Droplets were imaged under flow using fluorescence microscopy. A custom classification algorithm was applied to the Tau (Alexa Fluor 488) channel to assign each droplet as either homogeneous or as phase separated. This enabled reconstruction of phase diagrams with high resolution across a wide concentration space.

Native Tau demonstrated phase separated droplets at higher PEG8000 and protein concentrations in the phase diagram (Fig. 2). The phosphorylation state strongly modulated the phase behaviour in the phase diagrams. pTau^GSK3β^ displayed a reduced propensity for phase separation relative to pTau^ERK2^, consistent with its phosphorylation level and droplet characterization using fluorescence microscope (Fig. 1B). Across all conditions, oligomeric Bri2 BRICHOS exhibited a markedly inhibitory effect on Tau phase separation shifting the binodal boundary to higher protein and/or crowding concentrations (Fig. 2 and SI Fig. S5 and S6). To clearly address the effect of Bri2 BRICHOS itself, we also performed measurements with isolated Bri2 BRICHOS monomers. Similar to oligomeric species, monomeric Bri2 BRICHOS exhibited similar effects in the phase diagrams (SI Fig. S5 and S6). Together, these results demonstrate that both Bri2 BRICHOS and Tau phosphorylation pattern are key regulators of condensate formation, where phosphorylation promotes droplet formation while Bri2 BRICHOS acts as a potent suppressor of phase separation.

### Bri2 BRICHOS binds to pTau monomers in solution and in the droplet state

To study the molecular interactions between pTau monomers and Bri2 BRICHOS, solution NMR spectroscopy was applied to follow the intensity and chemical shift changes of the cross-peaks in the spectrum of monomeric isotopically ^15^N-labeled pTau upon addition of Bri2 BRICHOS. 2D ^1^H-^15^N-HSQC experiments were recorded on 100 μM ^15^N-labeled pTau in the absence and presence of 50 μM Bri2 BRICHOS. The spectra revealed Bri2 BRICHOS binding to both pTau^ERK2^ and pTau^GSK3β^, observed as a decrease of signal cross-peak intensity and chemical shift changes (Fig. 3). Notably, no new cross-peaks appeared upon Bri2 BRICHOS addition (Fig. 3). Some of the phosphorylated residues (pT175, pT181, pT231 and pT217) in pTau^ERK2^ are clearly visible and assigned in the ^1^H-^15^N-HSQC spectrum, exhibiting ^1^H chemical shifts of >9 ppm (Fig. 3A). In the presence of Bri2 BRICHOS, the signal intensities of these cross-peaks were drastically reduced, accompanied by chemical shift changes, in particular clearly visible for pT231 and pT217 (Fig. 3A and C).

**Figure 3.**
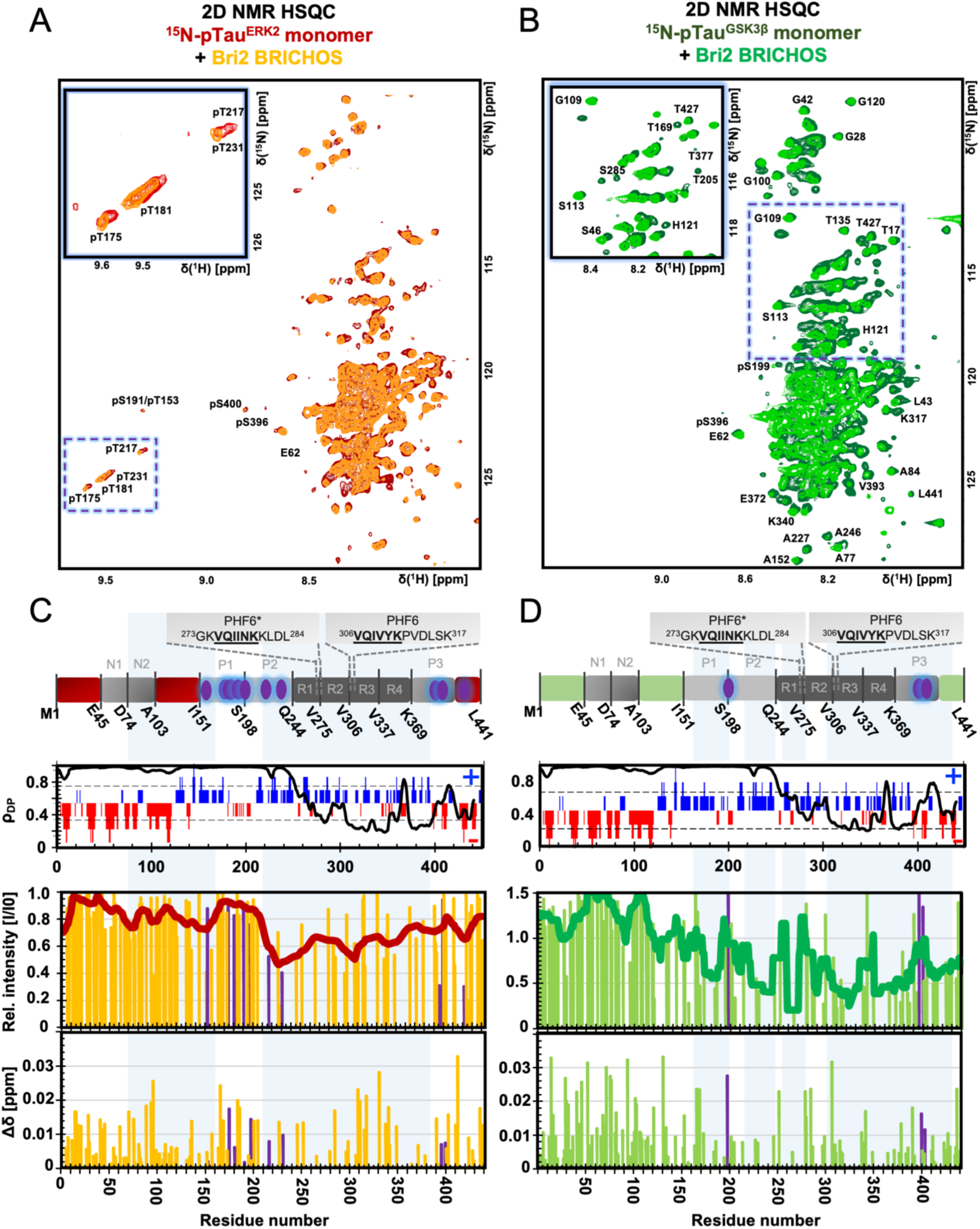
Bri2 BRICHOS binds to monomeric pTau. **(A, B)** ^1^H–^15^N-HSQC spectrum of 100 μM ^15^N-labeled monomeric **(A)** pTau^ERK2^ (red) and **(B)** pTau^GSK3β^ (dark green), and in the presence of 50 μM Bri2 BRICHOS (orange and light green, respectively). The NMR experiments were performed at 298 K in 20 mM HEPES buffer, pH 7.4. For pTau^ERK2^ several phosphorylated residues appear at resonances of >9 ppm in the proton dimension. **(C, D)** Top: Schematic view of pTau domains and molecular properties. Purple prolate spheroids represent confirmed phosphorylated residues. ρ_DP_ (residue-based droplet-promoting probabilities) and NCPR (net charge per residue) are calculated based on FuzDrop^53^ and CIDER ^54^ and plotted against the protein sequence. Bottom: Relative intensity (I/I0) and chemical shift changes (Δδ) determined from the spectra in (A) and (B). The solid line represents a smoothing function. Bars marked in purple represent phosphorylated residues.

Overall, the binding pattern of decreased signal intensity (up to around 50%) and chemical shift changes is more pronounced in the repeat domain, which includes the aggregation-prone hexapeptides PHF6/PHF6* motifs (Fig. 3), and in parts of the P1 and P2 regions and C-terminus for pTau^GSK3β^ (Fig. 3B and D). For pTau^ERK2^, interactions in the region covering both P2 and the repeat domain were observed (Fig. 3A and C). The P2 region of pTau^ERK2^ showed a clearly reduced signal intensity where two confirmed phosphorylated residues (pT217 and pT231) are present. The phosphorylated residues in pTau^ERK2^ in the sequence range of 170-200 did not show any decrease in signal intensity upon Bri2 BRICHOS binding, suggesting interactions with certain regions, i.e. the P1 & P2 regions and the repeat domain, including specific phosphorylated residues, such as pT217 and pT231. For pTau^GSK3β^, Bri2 BRICHOS binding affects similar regions as for pTau^ERK2^. In contrast to pTau^ERK2^, none of the phosphorylated residues of pTau^GSK3β^ exhibits a chemical shift at high ^1^H ppm values. No pronounced effects for two of the three confirmed phosphorylated residues of Bri2 BRICHOS could be observed, but one residue, pS199, displayed a pronounced chemical shift change.

We next set out to test the effect of Bri2 BRICHOS under conditions where pTau forms droplets. Addition of 8% PEG8000 induces phase separation and mimics the molecular wetting properties as an intracellular environment^55^. The presence of PEG8000 caused signal intensity reductions and chemical shift changes for some residues, whereas other residues were unaffected for both pTau^ERK2^ and pTau^GSK3β^ (Fig. 4). Remarkably, we observed that the phosphorylated residues at >9 ppm in pTau^ERK2^ were broadened and lost 50-90% of the original signal intensity in the spectrum (Fig. 4A). The strong effect of PEG8000 addition may point towards that the phosphorylated residues are involved in the phase-separated pTau system. Next, Bri2 BRICHOS was added to the pTau-PEG samples. Similar to the non-droplet state experiments (Fig. 3), Bri2 BRICHOS binding to pTau in the droplet state was evident (Fig. 4). Pronounced signal intensity reduction of 40-100% was apparent in the pTau^ERK2^ repeat domain and the P1/P2 regions (Fig. 4A, D). Interestingly, the effect in the droplet state was considerably larger compared to the ones in a non-droplet state. Apart from the relative intensity changes in the repeat domain, the P1/P2 regions were also strongly affected. Further, noticeable chemical shift changes in the regions of the N-terminus and P1/P2 regions were observed (Fig. 3C, 4D). Remarkably, the phosphorylated cross-peaks pT175 and pT181, which largely disappeared in the presence of PEG8000, reappeared upon addition of Bri2 BRICHOS to pTau^ERK2^ (Fig. 4A). This observation may suggest conformational changes of the protein induced by the Bri2 BRICHOS binding, resulting in a shift towards the monomeric and soluble pTau^ERK2^ state. In addition, new cross-peaks and/or large chemical shift changes were present. One of the cross-peaks, representing a phosphorylated residue (likely pT217 or pT231), clearly showed a chemical shift change upon addition of Bri2 BRICHOS (Fig. 4A). For pTau^GSK3β^, the binding pattern in the droplet state was remarkably similar to the dilute one (Fig. 3), but with more pronounced effects in the repeat region of residues 270-305 including the hexapeptide PHF6* (Fig. 4E).

**Figure 4.**
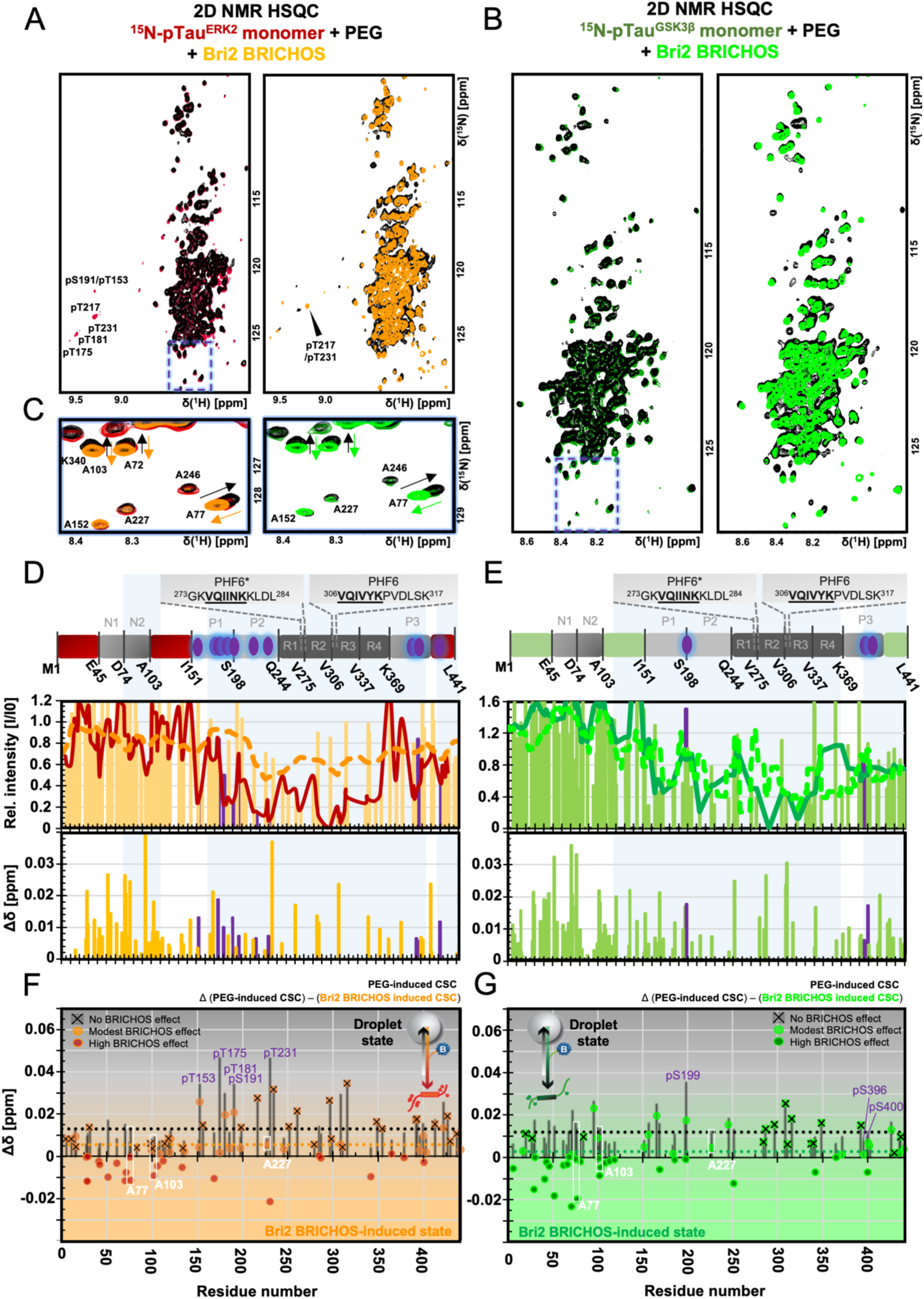
Bri2 BRICHOS interacts with pTau in the droplet state. **(A, B)** ^1^H–^15^N-HSQC NMR spectrum of 100 μM ^15^N-labeled monomeric **(A)** pTau^ERK2^ (red) and **(B)** pTau^GSK3β^ (dark green), and in the presence of 8% PEG8000 (black) and 50 μM oligomeric Bri2 BRICHOS (orange and light green, respectively). The NMR experiments were performed at 298 K in 20 mM NaP buffer, pH 7.4. For pTau^ERK2^ several phosphorylated residues appear at >9 ppm in the proton dimension. Zoom-in images from (A, B) is presented in **(C)**. **(D, E)** Top: Schematic view of pTau. Purple prolate spheroids represent confirmed phosphorylated residues. Bottom: Relative intensity (I/I0) and chemical shift changes (Δδ) determined from the spectra in (A) and (B). I/I0 is the relative NMR signal intensity of ^15^N-pTau_441_ in 8% PEG8000 with and without Bri2 BRICHOS. The solid line represents a smoothing function. The dashed line represents the effect of Bri2 BRICHOS on pTau in the non-droplet state from Figure 3. Bars marked in purple represent confirmed phosphorylated residues. **(F, G)** Δδ induced by PEG8000 (black bars) compared to the effect of Bri2 BRICHOS on pTau^ERK2^ (red) and pTau^GSK3β^ (green). The Δδ effects from Bri2 BRICHOS from (D, E) were subtracted from the PEG8000 effects (black bars) to visualize the opposing regulatory effect of Bri2 BRICHOS on the droplet state (circles). The dashed lines represent the average values from PEG8000-respectively Bri2 BRICHOS effects, where the value for the latter is approaching zero, corresponding to a shift back to the reference cross-peak position of pTau alone.

In the presence of PEG8000, chemical shift changes for pTau^ERK2^ and pTau^GSK3β^ were uniformly distributed throughout the protein with more pronounced changes in the phosphorylated regions. Addition of Bri2 BRICHOS seemingly counteracts these chemical shift changes back to the reference state of pTau alone (Fig. 4A-C). To further expand and quantify these observations, the chemical shift changes from PEG8000 versus Bri2 BRICHOS addition were plotted together (Fig. 4F, G and SI Figs. S7, S8). A comparison between the chemical shift changes from the PEG8000-induced droplet-state, versus the effect from Bri2 BRICHOS, revealed counteracting effects for cross-peaks initially experiencing PEG8000-induced effects (62% of the pTau^ERK2^ cross-peaks, 75% of the pTau^GSK3β^ cross-peaks). Some cross-peaks were shifted back to the exact original position (represented by a difference of zero), and other cross-peaks, illustrated by examples of A77 and A103, had even larger effects by Bri2 BRICHOS, resulting in a cross-peak position that passed the reference point (which is represented by negative values in the difference plot). A third set of peaks basically did not experience any effect by Bri2 BRICHOS.

The chemical shifts of the cross-peaks upon the addition of Bri2 BRICHOS resembled the ones without a PEG8000-induced droplet state. Therefore, we compared the chemical shifts changes from Bri2 BRICHOS in a droplet state and from the dilute state (SI Figs. S7, S8). Surprisingly, the chemical shift changes from both states agreed to a high extent, suggesting a similar Bri2-BRICHOS-pTau bound state regardless of whether pTau previously was in a condensed- or in the dilute state. Based on these observations, together with the Bri2 BRICHOS inhibitory effect on condensates, we conclude that Bri2 BRICHOS exhibits an opposing regulatory effect of Tau condensates.

Taken together, beyond differences in the phosphorylation pattern, Bri2 BRICHOS binds to pTau^GSK3β^ and pTau^ERK2^ in both non-droplet and droplet state conditions. The binding regions are dependent on the local environment (non-droplet or droplet state) and phosphorylation pattern. The MTB region of Tau is clearly affected by Bri2 BRICHOS, irrespectively of the phosphorylation pattern or condensation state. In the non-droplet state, both pTau variants further exhibit affected P2 regions and for pTau^GSK3β^ also the P1 region is modulated. Interestingly, for the droplet state condition, both P1 and P2 regions are involved in the Bri2 BRICHOS interactions for both pTau variants (Fig. 4). The interactions between Bri2 BRICHOS and pTau variants are significantly stronger compared to the ones for native Tau^41^, and exhibit opposing regulatory effects on the condensate state towards a shift to a non-droplet and soluble state.

## DISCUSSION

The balance between physiological and pathological Tau states is tightly associated with hyperphosphorylation and proteostasis mechanisms^13^. However, ways to prevent pathological triggers have remained elusive. The molecular chaperone system is one component of proteostasis to regulate and counteract protein misfolding. Here, we show that Tau phosphorylation promotes Tau condensation and modulates the fibrillation behavior, whereas chaperone-mediated regulation by Bri2 BRICHOS counteracts phosphorylation-induced phase separation and generally inhibits Tau fibrillation with different phosphorylation patterns.

The droplet formation propensity of Tau is dependent on the number of phosphorylated residues, where each phosphorylated residue contributes with −1 to the overall net charge. Electrostatics are important for Tau phase separation, in particular in specific regions essential for interactions involved in phase separation^16,17^. pTau^ERK2^ is the most phosphorylated variant, thus exhibiting the greatest negative net charge in our study (SI Fig. 1), demonstrated the highest droplet propensity (Figs. 1C, D, and 2). Modest phosphorylation of the pTau^GSK3β^ variant (with a net charge around zero) showed reduced condensate propensity compared to pTau^ERK2^ (Figs. 1C, D and 2), while native Tau (with a positive net charge), exhibited the lowest droplet propensity.

While the condensation propensity is seemingly largely governed by the overall net charge, the aggregation propensity likely is more dependent on the charge distribution and phosphorylation pattern of specific regions, where pTau^GSK3β^ was most aggregation-prone followed by native Tau and pTau^ERK2^ (Fig. 1B). The equilibrium from the droplet state may have been shifted towards fibril formation, strengthened by a net charge close to zero at physiological pH providing favorable hydrophobic interactions necessary for amyloid formation, in comparison to electrostatic interactions necessary for droplet formation^8,10,11,49,50^. Several phosphorylated sites are located outside the fibril core, suggesting that electrostatic repulsion between monomers and/or the fuzzy fibril coat may protect against nuclei formation. Hence, the phosphorylation pattern affects droplet formation and fibrillation differently.

The condensate propensity for all Tau variants was reduced by Bri2 BRICHOS, along with a general delay of the fibrillation kinetics (Figs. 1 and 2). The fibril inhibition was most pronounced for the aggregation-prone pTau^GSK3β^ (Fig. 1B). The ability of Bri2 BRICHOS to inhibit Tau aggregation regardless of the phosphorylation pattern suggests similar underlying inhibitory mechanisms dominated by suppression of surface-catalyzed secondary nucleation processes, as previously reported for native Tau and Bri2 BRICHOS^41^.

Phase boundaries were shifted to a higher extent for both pTau variants in the presence of Bri2 BRICHOS compared to native Tau (Fig. 2 and SI Fig. S6). Further, Bri2 BRICHOS at equimolar concentrations reduced the formation of droplets, counteracted droplet maturation, and promoted a more liquid-like state over time (Figs. 1, 4 and SI Figs. S2-S4). Following the theory of a condensate state as a prerequisite or intermediate for nucleation, Bri2 BRICHOS may inhibit such early aggregation precursors in pTau condensates. Notably, inhibition of Tau-associated neurotoxicity effect by Bri2 BRICHOS has previously been shown^41^.

Incorporation of Bri2 BRICHOS into pTau condensates is indicative of specific interactions between the two proteins (Fig.1), where we determined Bri2 BRICHOS binding sites using solution NMR. We found that Bri2 BRICHOS binds to monomeric pTau in solution and to the pTau droplet state (Figs. 3 and 4). Similar to native Tau^41^, the interactions between pTau and Bri2 BRICHOS are more pronounced within the condensate state, with the P1/P2 and MTB region especially affected, including regions with phosphorylated residues (Figs. 3 and 4). The MTB region exhibits a positive net charge and is sparsely phosphorylated in our and other studies^23^, including disease-related Tau protein deposits found in brain^5^. Bri2 BRICHOS is negatively charged, which suggests contributions from electrostatic interactions. The projection domain (P1/P2) is important for binding towards microtubules and for long-range interactions within the protein to the MTB region, which is overall important for fibril formation^46^. Bri2 BRICHOS binding to these regions may hence modulate or partly block interaction sites, which are crucial for droplet and fibril formation.

Our NMR experiments revealed that Bri2 BRICHOS had a counteracting effect on PEG-induced chemical shift changes (Fig. 4F, G), indicating a stabilization of a state resembling the monomeric state in solution (Fig. 3, SI Figs. S7, S8). Remarkably, residues of pT217 and pT231 in pTau^ERK2^ are strongly affected by Bri2 BRICHOS (Fig. 3 and 4). These residues may be directly involved in the binding of Bri2 BRICHOS, or in the proximity of the binding sites involving additional residues. pT231 and pT217 have been shown to act as early clinical biomarkers for AD^19^, and the ability of Bri2 BRICHOS to interact with these residues may shed light on important molecular events in disease initiation and/or progression.

Taken together, this study provides mechanistic insights into the multifaceted behavior of disease-related phosphorylated Tau and how its condensation and fibrillation can be opposingly regulated by the phosphorylation pattern and the chaperone-like Bri2 BRICHOS domain. These findings can be rationalized into a model of mechanism-of-action, describing how phosphorylation and chaperone interactions regulate Tau phase separation and fibril formation (Fig. 5). Extensive phosphorylation promotes biomolecular condensate formation, which can be counteracted by Bri2 BRICHOS interactions in the dilute phase and droplet state. Bri2 BRICHOS is integrated into the droplets, which seemingly prevents maturation of the droplets – a state that potentially represents an early precursor of nucleation and fibril formation. Eventually, Bri2 BRICHOS generally inhibits Tau aggregation for both investigated phosphorylation states, suggesting a genetic inhibitory effect that is not perturbated by the phosphorylation pattern.

**Figure 5.**
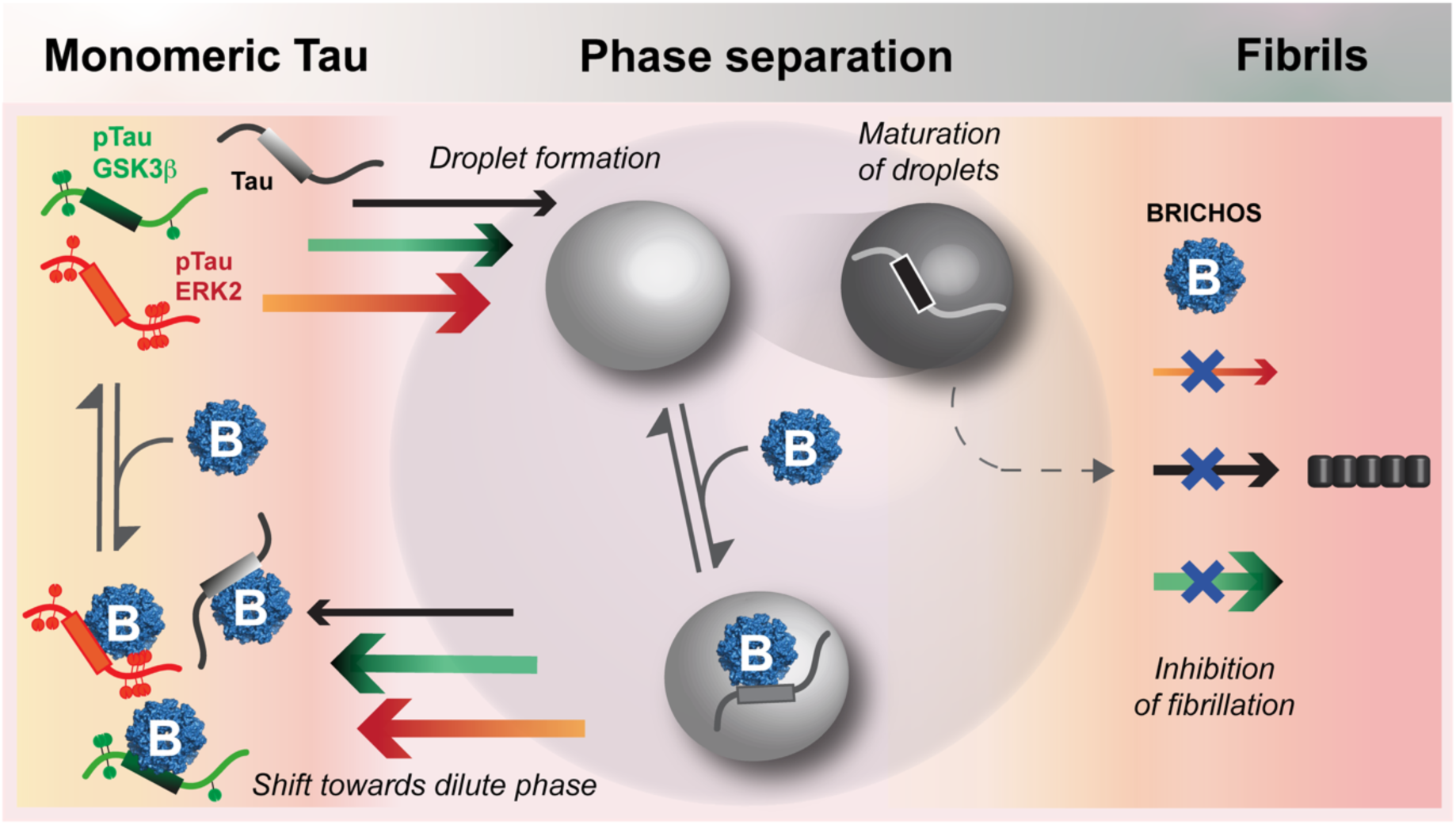
Mechanistic model of how the chaperone-like Bri2 BRICHOS domain counteracts phosphorylation-promoted Tau phase separation and fibrillation. Phosphorylation of Tau proteins promotes phase separation, where pTau^ERK2^, the most charged one, exhibited the highest propensity to form droplets, followed by pTau^GSK3β^ and native Tau. Bri2 BRICHOS binds to all Tau variants in the dilute and droplet state, redirecting Tau from condensation towards the dilute phase. While the different Tau variants exhibit different aggregation propensities (indicated by the thickness of the arrows), Bri2 BRICHOS generally inhibits Tau fibrillation irrespectively of the phosphorylation patterns. The Bri2 BRICHOS structure is based on PDB ID 8RNU^56^.

The chaperone-like 14-3-3ζ protein has also been shown to inhibit fibrillation of pTau^GSK3β^ via interactions with P1/P2 and the MTB region^57^, as well as a reduction of pTau aggregation in neurons and promotion of condensates at sub-stoichiometric concentrations^58^. Another example of monomeric Tau binding and subsequent effects on fibrillation and solubility is the immunophilin FKBP12^59^. The MTB region binds to the negatively charged microtubules in cells and contains the aggregation-prone hexapeptides PHF6/PHF6*. This MTB region is vital for amyloid fibril formation and is also part of the fibril core^60–62^. Several anti-amyloidogenic molecules have shown a preferred binding to this region, to sterically hinder the aggregation-prone regions to interact and avoid further nucleation/polymerization^63^. Hence, Bri2 BRICHOS binds to the same region in pTau and native Tau^41^, highlighting the crucial role of this region.

The interaction between Bri2 BRICHOS and pTau raises the question whether the observed effects of Bri2 BRICHOS can be related to its physiological function. The BRICHOS domain is cleaved and secreted to the extracellular space after Bri2 processing, and Tau is mainly associated to microtubules in neurons, or in disease, as aggregated species in the soma. However, Tau spreads from cell to cell, possibly by both active and passive transport^64^, which provides possibility for extracellular interactions with Bri2 BRICHOS. Further, Bri2 BRICHOS is also taken up by cells^64,65^ and may hence interact with misfolded species intracellularly. Based on this background, future research may concern strategies for external administration or modulation of endogenous expression of Bri2 BRICHOS or other chaperone-like proteins as potential therapeutics.

## CONCLUSION

This study demonstrates that the extent of phosphorylation governs the biomolecular condensation and fibrillation behavior of Tau and that these effects can be counteracted by the Bri2 BRICHOS domain. These findings enhance our molecular understanding of how disease-related processes can be regulated and safeguarded by chaperone-like proteins.

## MATERIAL AND METHODS

### Recombinant proteins

Recombinant full-length Tau_1-441_ protein and the Bri2 BRICHOS domain were expressed and purified following previously described protocols^39,41,48,66^. In short, Tau was expressed in BL21(DE3) *E. coli* cells and induced for 2 hours by addition of 0.4 mM IPTG at 37 °C. Cell pellets were purified by resuspension in 50 mM NaP buffer pH 6.5, 1 mM EDTA, and one protease inhibitor cocktail tablet 1x (cOmplete, EDTA-free, Roche). The cells were lysed by sonication, followed by centrifugation to remove larger cell parts. The supernatant was heat-treated and centrifuged to remove proteins from the heat-stable Tau protein solution. The supernatant containing the remaining soluble proteins including Tau was kept at −20 °C before being thawed for further purification. Tau was separated from other remaining proteins by cation-exchange chromatography (HiTrap SP HP, Cytiva), using an increasing NaCl concentration gradient for elution. The fractions were analyzed on SDS-PAGE, and the fractions containing pure Tau were pooled together. The excess NaCl was removed using a HiPrep™ 26/10 Desalting column (Cytiva) into 20 mM NaPi pH 7.4 with 0.2 mM EDTA or 20 mM HEPES pH 7.4 or 8, and aliquoted and stored at −20 °C. An extinction coefficient of 7550 M^−1^ cm^−1^ was used to determine the Tau concentration. ^15^N-labeled Tau was expressed and purified using similar methods, but using minimal M9 media instead of LB media, and supplement of ^15^NH_4_Cl, for the expression. All buffers and solvents used for the experiments were filtered and degassed before usage.

The Bri2 BRICHOS domain in monomeric and oligomeric assembly states, constituting residues 113-231 of the Bri2 protein with a R221E mutation and two additional N-terminal residues (with the sequenc: GSQTI EENIK IFEEE EVEFI SVPVP EFADS DPANI VHDFN KKLTA YLDLN LDKCY VIPLN TSIVM PPRNL LELLI NIKAG TYLPQ SYLIH EHMVI TDRIE NIDHL GFFIY ELCHD KETYKL), were expressed within the plasmid pT7NT*-Bri2 113-231 R221E (Addgene plasmid # 138134) in SHuffle T7 (K12) *E. coli* cells according to published work^39^. NT* is used as a solubility tag during expression and purification. Briefly, expression was performed at 20 °C overnight after induction by addition of 0.5 mM IPTG. The cells were harvested by centrifugation and resuspended in 20 mM Tris-HCl pH 8, followed by sonication, centrifugation, and loading of supernatant onto a Ni-NTA column (GE Healthcare Bio-Science) and elution with imidazole. Imidazole removal was performed by dialysis in 20 mM Tris pH 8 together with addition of thrombin to remove the N-terminal solubility tag (His6-NT*) overnight. The His6-NT* tag was separated from Bri2 BRICHOS monomers and oligomers by a Ni-NTA gravity column. Bri2 BRICHOS was collected through size exclusion chromatography (SEC), fractions analyzed by SDS-PAGE, and the concentrations were determined with an extinction coefficient of 9065 M^−1^ cm^−1^.

### *In vitro* phosphorylation of Tau

Native Tau and ^15^N-Tau were phosphorylated *in vitro* by two different kinases following published protocols in literature^23,66^. Activated GSK3β (Sigma-Aldrich SRP0356) at 0.18 μM was mixed to 120 μM Tau, with 2 mM ATP, 2 mM TCEP, 5 mM EDTA, 1x protease inhibitor cocktail tablet (cOmplete, EDTA-free, Roche), and 5 mM MgCl_2_ in 20 mM HEPES pH 7.4 buffer and incubated for 15 h at 30 °C while shaking at 300 rpm. For ERK2 (Sigma-Aldrich 14-550) phosphorylation, 0.1 μM activated kinase was used for 130 μM Tau and supplemented with 2.5 mM ATP, 1 mM TCEP, 2 mM EDTA, 50 mM NaCl, 1x protease inhibitor cocktail tablet (cOmplete, EDTA-free, Roche), and 12 mM MgCl_2_ in 40 mM HEPES pH 8 buffer and incubated 3 h at 37 °C. After the phosphorylation reaction, the enzyme was deactivated and precipitated by heat-treatment and 60 min centrifugation at 4 °C. The supernatant was kept and buffer exchanged to 20 mM NaP pH 7.4. The quality of the obtained phosphorylated Tau variants was analyzed by SDS-PAGE and NMR spectroscopy.

### Fluorescence microscopy

Fluorescence microscopy imaging was performed using a Nikon Ti2-AX inverted microscope (Nikon) with a resonant scanner and 2x tunable GaAsP detectors. Samples of 15 µM pTau and 8% (w/vol) PEG8000 with 1% Atto-488 fluorophore in 30 µl were imaged in µ-Plate 96 Well Square (Ibidi) after 45-min equilibration at room temperature in 20 mM sodium phosphate, 0.2 mM EDTA pH 7.4 using a 40x objective (Nikon). Visualization of Bri2 BRICHOS incorporation into pTau droplets was performed with 1% Bri2 BRICHOS-Atto-655. Fluorescent labeling of Tau and Bri2 BRICHOS was obtained with amine-reactive Atto-488 NHS-ester dye (Sigma-Aldrich, 41698) and Atto-655 NHS-ester dye (Sigma-Aldrich, 76245) previously described in detail^41^.

Dilution experiments for viscosity characterization of the samples were performed with 15 µM pTau^ERK2^ and pTau^GSK3^ labeled with Atto-488 in the absence or presence of 3 µM Bri2 BRICHOS-Atto-655. Identical samples were used for 50% dilution, sequentially imaged as described above.

### Thioflavin T (ThT) fluorescence kinetics

The fibrillation of pTau in the absence and presence of Bri2 BRICHOS was followed by a Thioflavin T (ThT) assay. 20 µM pTau in 20 mM NaP pH 7.4 and 0.2 mM EDTA supplemented with 50 µM ThT (λ_ex_: 440 nm, λ_em_: 480 nm) was used in 384-well plates (Corning), 20 µl per well with 6 replicates, and fluorescence measured every 10 minutes in a FLUOStar Omega plate reader (BMG Labtech) at 37 °C. Orbital shaking at 300 rpm for 9 minutes between the fluorescence reading cycles and two glass beads with a diameter of 1 mm per well were used to induce fibrillation without co-factors. The assay was repeated several times. The ThT kinetic curves were normalized to the plateau of each condition without Bri2 BRICHOS and fitted to a sigmoidal curve to obtain phenomenological parameters such as the aggregation halftime, t_1/2_, where half of the monomers are depleted.

### NMR spectroscopy

2D NMR ^1^H-^15^N-HSQC experiments were performed on a 700 MHz Bruker Avance spectrometer equipped with a cryogenic probe. 50 µM unlabeled oligomeric Bri2 BRICHOS was added upon 100 µM monomeric ^15^N-pTau in 20 mM HEPES or NaP buffer pH 7.4 and 0.2 mM EDTA. Additional experiments were performed with 8% (w/vol) PEG8000 for phase separation conditions. The experiments were performed at 298 K with 90%/10% H_2_O/D_2_O and the experimental temperature was calibrated with an external thermometer. The spectra with and without Bri2 BRICHOS, both in aqueous solution and during phase separation conditions were compared by calculating the relative intensities based on the amplitude intensities and determination of the chemical shift changes (csc). The csc were calculated from Equation 1:

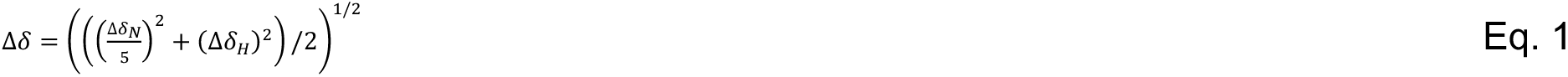

The cross-peak assignment of pTau (BMRB ID 50701) was achieved by a comparison to published work ^41,67–74^. Data were processed and analyzed with Topspin v.4.2.0 (Bruker) and Poky^75^ (University of Colorado Denver).

### Fabrication of microfluidic devices

Microfluidic devices were fabricated following established protocols^51,52^. Device layouts were designed using AutoCAD and fabricated via standard soft photolithography. Briefly, SU-8 3050 photoresist (A-Gas Electronic Materials Limited) was spin-coated onto polished silicon wafers (MicroChemicals GmbH) to achieve a thickness of ∼50 µm. The coated wafers were soft-baked, exposed to UV light through a photomask, post-baked, and developed in propylene glycol monomethyl ether acetate (PGMEA; Sigma-Aldrich) to produce the master mould. Polydimethylsiloxane (PDMS; Sylgard 184, Dow Corning; 10:1 base-to-crosslinker ratio) was poured over the master, cured at 60 °C for 2 h, and subsequently cut to size. Inlet and outlet ports were introduced by punching. Following sonication in isopropanol, PDMS slabs were bonded to glass slides (Epredia) using oxygen plasma treatment (60% power, 30 s; Diener Femto). To render channel surfaces hydrophobic, devices were treated with 1% (v/v) trichloro(^1^H,^1^H,^2^H,^2^H-perfluorooctyl)silane (Sigma-Aldrich) dissolved in HFE-7500 (3M™ Novec™ engineered fluid), and then baked at 95 °C prior to use.

### Generation of phase diagrams

Multidimensional phase diagrams were generated using PhaseScan, a semi-automated combinatorial droplet microfluidics platform, following established approaches^51,52^. In each experiment, aqueous input streams comprising pTau or Tau, monomeric or oligomeric Bri2 BRICHOS, buffer, and PEG8000 were combined within a droplet-generating microfluidic device. Flow rates of the aqueous solutions and the continuous oil phase were precisely controlled using pressure-driven Flow EZ™ systems (Fluigent). Concentration space was sampled by dynamically varying the relative flow rates of the aqueous inputs, enabling the generation of droplets spanning a continuous range of component concentrations. To obtain quantitative readout of composition, each aqueous component was fluorescently barcoded. pTau and Tau were labeled with Alexa Fluor 488 (1% NHS labeling), Bri2 BRICHOS (monomeric or oligomeric) with Alexa Fluor 647 (1% NHS labeling), and PEG8000 with Alexa Fluor 546 (Thermo Fisher Scientific). Droplets were formed in HFE-7500 oil supplemented with 1.2% (w/v) fluoro surfactant (RAN Biotechnologies), which was introduced at a constant flow rate. Droplet populations were imaged on-chip under continuous flow using an epifluorescence microscope (Cairn Research) equipped with a 10× objective. Subsequent image processing was carried out using custom Python-based analysis pipelines. Individual droplets were segmented and subjected to quality control filtering, after which fluorescence intensities corresponding to each barcode were extracted, normalized to droplet volume, and converted to absolute concentrations using calibration curves. Each droplet was then classified as either homogeneous or phase-separated based on the presence of condensates in the Tau channel. The resulting datasets were used to reconstruct high-resolution phase diagrams, which were visualized as averaged scatter plots representing the distribution of phase behaviour across the sampled concentration space.

## Code availability

The PhaseScan analysis code used in this study is available at GitHub (https://github.com/rqi14/PhaseScan).

## Acknowledgements

This study was supported by the Swedish Research Council (AA), the Swedish Alzheimer Foundation (AA), the Swedish Brain Foundation (AA), CIMED (AA), Loo and Hans Osterman Foundation (CM), Foundation for Geriatric Diseases at KI (CM), Åhlen Foundation (CM, AA), Magnus Bergvall Foundation (CM), David and Astrid Hagelén Foundation (CM), Gun and Bertil Stohnes Foundation (CM), and the Stiftelsen för Gamla Tjänarinnor (CM), the Petrus and Augusta Hedlunds Stiftelse (CM) and the European Research Council (ERC) [DiProPhys 101001615] (TA and TPJK). This study was in part performed at the Live Cell Imaging core facility/Nikon Center of Excellence, at Karolinska Institutet, Sweden, supported by the KI infrastructure board (1–48/2024) and the Olle Engvist Foundation (235-0496).

## Notes

The authors declare no competing financial interest.

