## Supporting Information for "Opposing Regulation of Tau Condensation by Phosphorylation Levels and Molecular Chaperone Interactions"

**TABLE OF CONTENT**

**Supporting Figures**……………………………………………………………………………………

**SI Figure S1.** Schematic presentation of model system using three different Tau variants and Bri2 BRICHOS.

**SI Figure S2.** Bri2 BRICHOS inhibits pTau droplet formation.

**SI Figure S3.** Bri2 BRICHOS and pTau droplets incubated for 24 hours.

**SI Figure S4.** Dilution of mature droplets with and without Bri2 BRICHOS.

**SI Figure S5.** High-resolution phase diagrams of Tau and pTau in the presence of monomeric and oligomeric Bri2 BRICHOS.

**SI Figure S6.** Binodal phase boundaries determined by PhaseScan for the different Tau variants.

**SI Figure S7.** Bri2 BRICHOS counteracts PEG8000-induced chemical shift changes in $pTau^{ERK2}$.

**SI Figure S8.** Bri2 BRICHOS counteracts PEG8000-induced chemical shift changes in $pTau^{GSK3\beta}$.

**References**…………..……………………………………………………………………………

| | | Net charge*<br>pI*<br>Monomer, pH 7.4 | Confirmed phos. residues (by NMR spectroscopy) | Confirmed phos. residues in literature | |
|---|---|---|---|---|---|
| 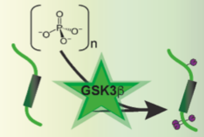  | pTau GSK3β | -3.84<br>6.69 | pS199<br>pS396<br>pS400 | **Chakraborty et al.**<br>pT175<br>pT181<br>pT205<br>pT231<br>pS404 | **Kučinskas et al.**<br>pS199<br>pT205<br>pS396<br>pS400<br>pS404 |
| 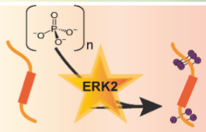  | pTau ERK2 | -12.64<br>6.055 | pT153<br>pT175<br>pT181<br>pS191<br>pS198<br>pT217<br>pT231<br>pS396<br>pS400<br>pS422 | **Chakraborty et al.**<br>pT175<br>pT181<br>pS202<br>pT205<br>pS210 | pT212<br>pS235<br>pT245<br>pS396<br>pT403<br>pS404 |
| 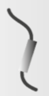 | Tau | 1.44<br>7.94 | 0 | 0 | |
| 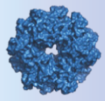 | BRICHOS | -12.48<br>4.6 | 0 | 0 | |

* based on phosphorylation pattern determined by NMR spectroscopy

**SI Figure S1. Schematic presentation of model system using three different Tau variants and Bri2 BRICHOS.** The phosphorylation patterns were determined by NMR in this study. Net charge and pI were calculated with ExPASy[1], using the phosphorylation sites determined by NMR. Phosphorylation patterns reported in literature are presented in right panel[2,3]. The Bri2 BRICHOS illustration (blue) is based on PDB ID 8RNU[4].

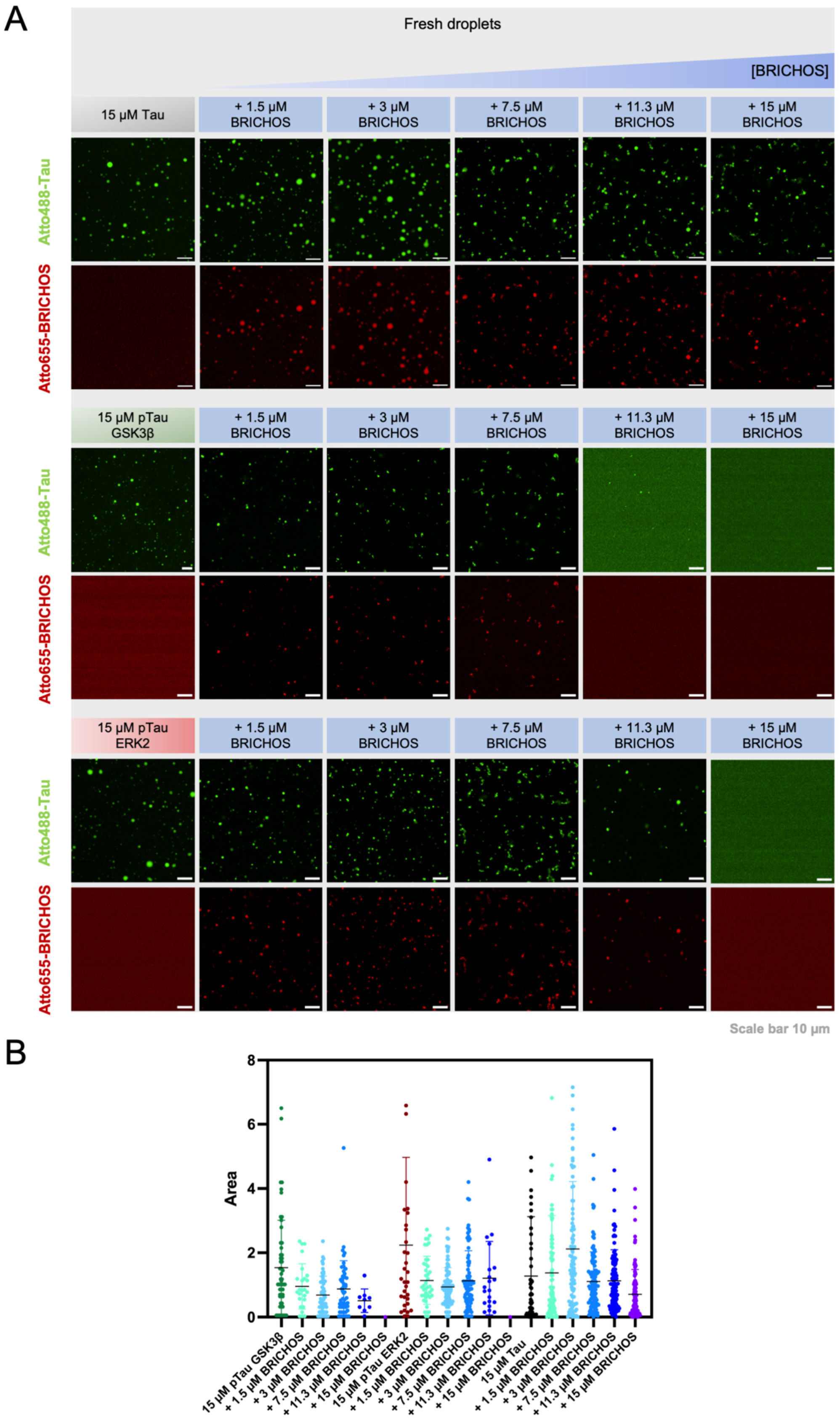


**SI Figure S2. Bri2 BRICHOS inhibits pTau droplet formation**. Whole concentration series of the droplets shown in Fig. 1A in the main manuscript. Fluorescence microscopy images of 15 µM non-phosphorylated, native Tau, or pTau with 0, 1.5, 3, 7.5, 11.3, and 15 µM Bri2

BRICHOS concentrations in droplet-promoting buffer of 8% PEG8000 and 20 mM NaP pH 7.4. 1% Atto-655 Bri2 BRICHOS and/or 1% Atto-488 Tau were supplemented to the samples to visualize the presence or absence of droplets with florescence. The scale bar represents 10 µm.

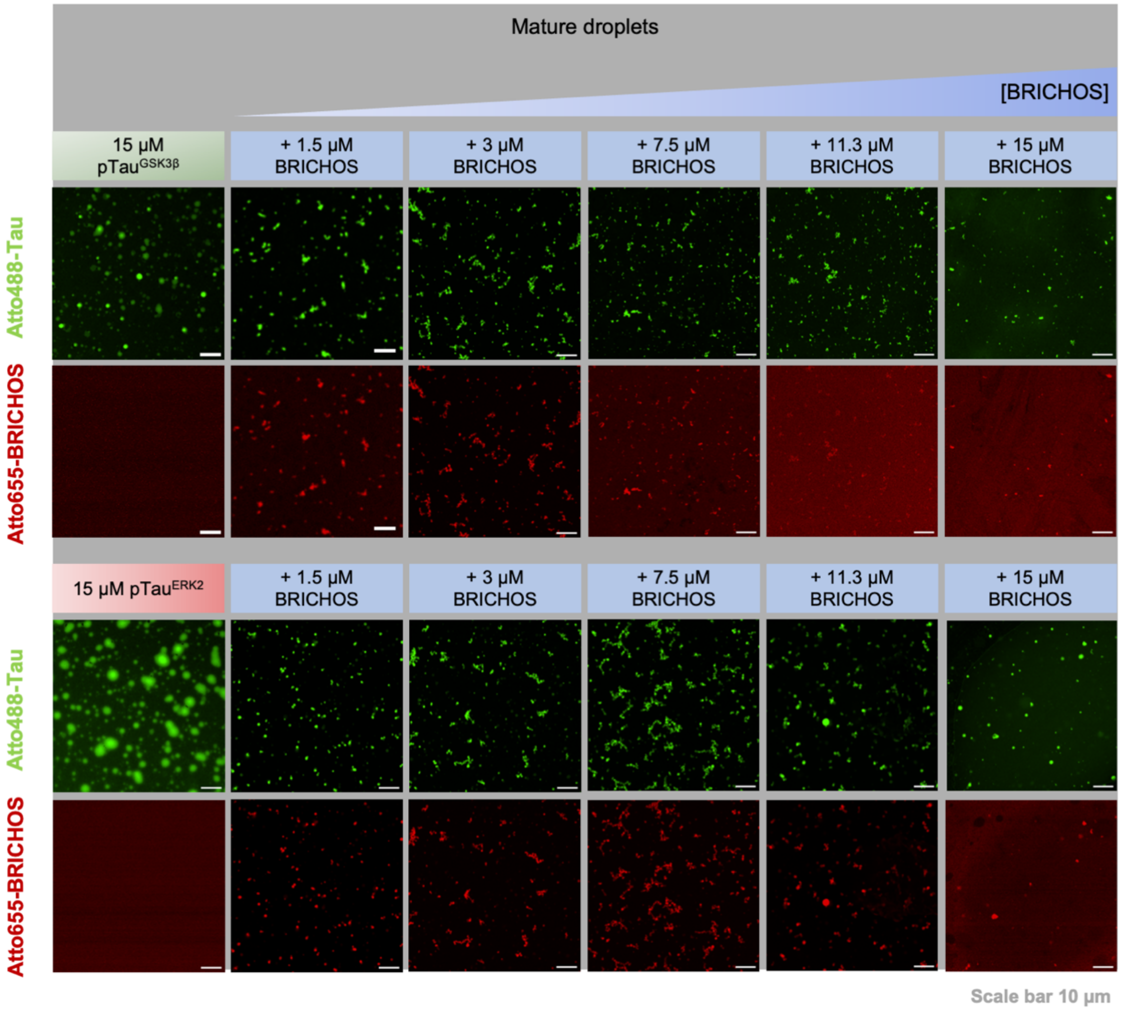


**SI Figure S3. Bri2 BRICHOS and pTau droplets incubated for 24 hours.** Whole concentration series of the droplets shown in Fig. 1C and SI Fig. S2 after 24 h incubation. The scale bar represents 10 µm.

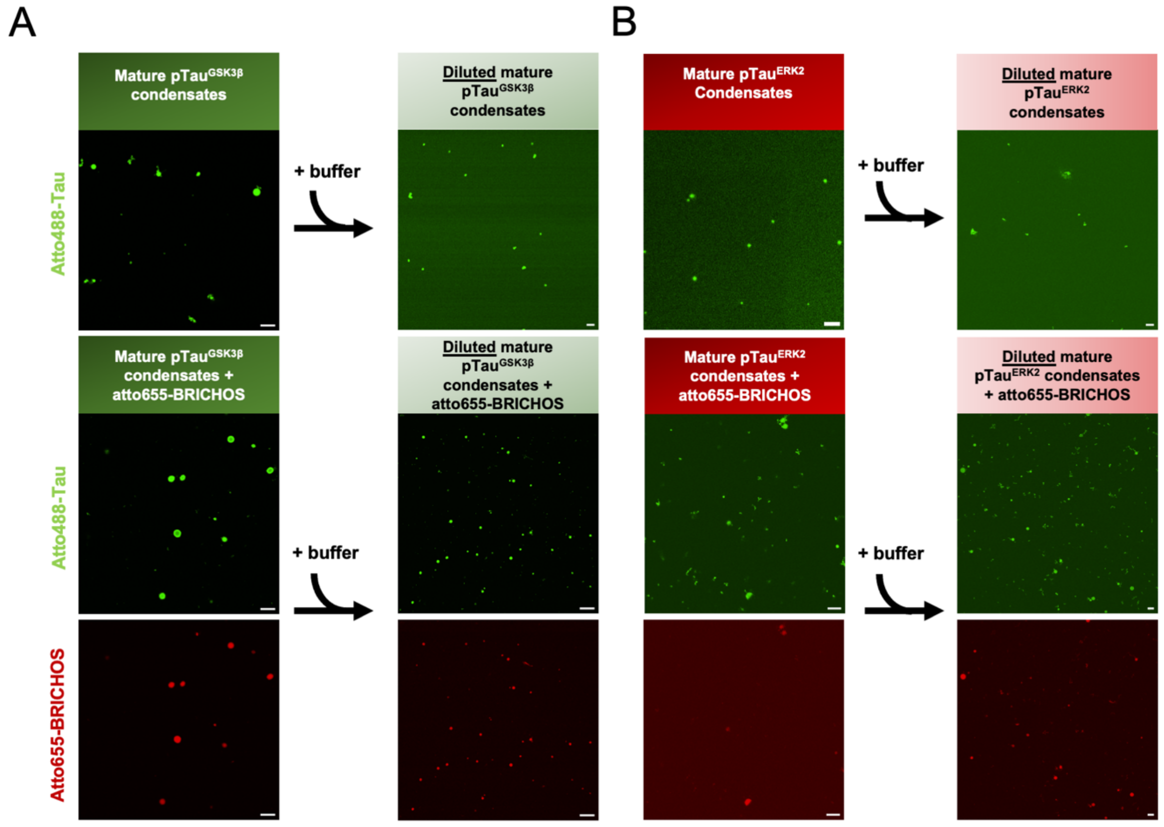


**SI Figure S4. Dilution of mature droplets with and without Bri2 BRICHOS.** 24 h mature droplets of 15 µM pTau in the absence or presence of 3 µM Bri2 BRICHOS, diluted 50% with buffer. The scale bar represents 10 µm.

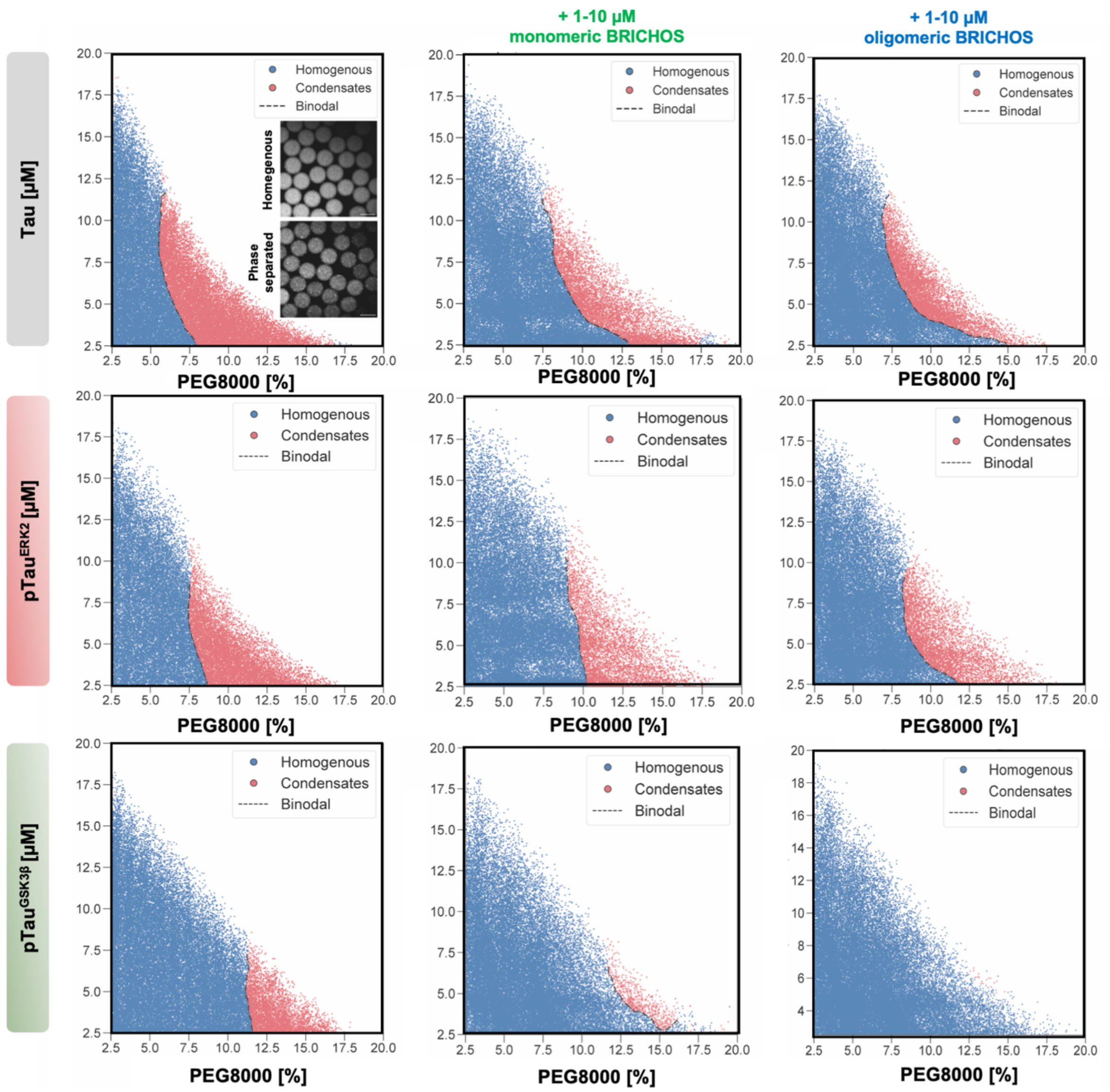


**SI Figure S5. High-resolution phase diagrams of Tau and pTau in the presence of monomeric and oligomeric Bri2 BRICHOS.** High-resolution phase diagrams generated using PhaseScan for Tau, pTau$^{ERK2}$, and pTau$^{GSK3\beta}$ as a function of PEG8000 concentration, and the effect of monomeric (middle panel) and oligomeric (right panel) Bri2 BRICHOS. Each point represents an individual microdroplet, classified as homogeneous (blue) or phase-separated (red). Dashed lines indicate the inferred binodal phase boundaries. Extracted phase boundaries are presented in SI Figure S6. The number of analyzed microdroplets per condition from left to right, top to bottom, is: n = 44994, n = 46641, n = 49398, n = 36522, n = 39002, n = 44052, n = 50055, n = 43590, n = 51541. Representative widefield images are displayed; scale bar 100 µm.

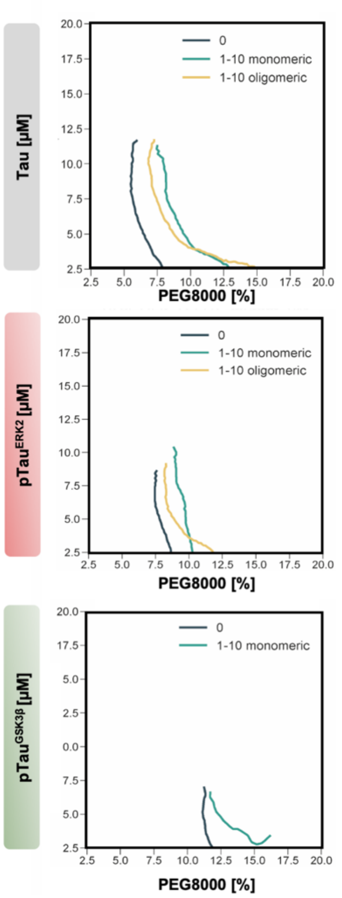


**SI Figure S6. Binodal phase boundaries determined by PhaseScan for the different Tau variants.**

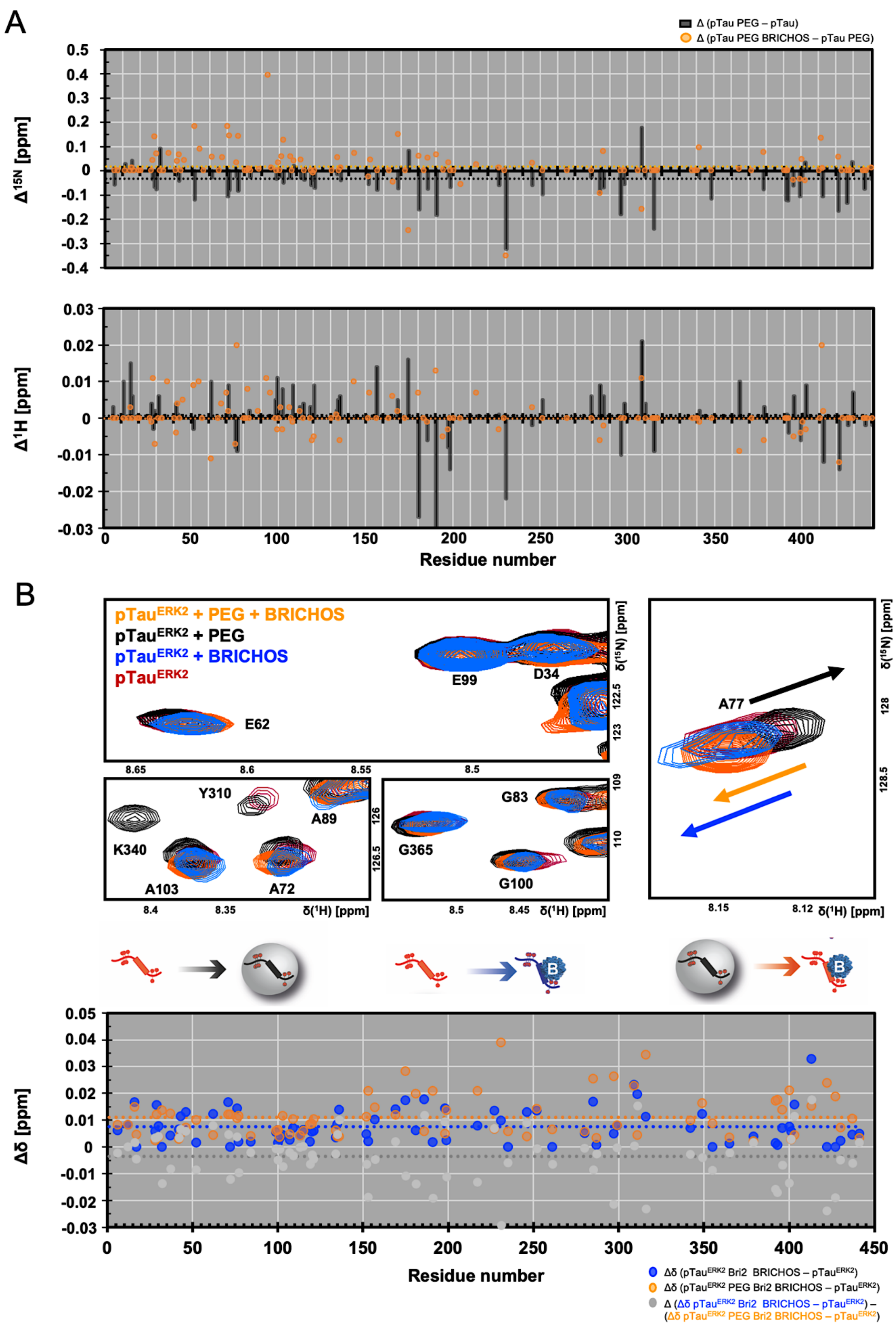


**SI Figure S7. Bri2 BRICHOS counteracts PEG8000-induced chemical shift changes in pTau$^{ERK2}$. (A)** $\Delta^{15}$N and $\Delta^{1}$H chemical shifts of pTau$^{ERK2}$ in the presence of 8% PEG8000 (black bars) or the Bri2 BRICHOS effects on pTau$^{ERK2}$ in PEG8000 (orange circles). **(B)** $^{1}$H–$^{15}$N-HSQC

NMR spectra of 100 μM $^{15}$N-labeled monomeric pTau$^{ERK2}$ (red) in the presence of 50 μM oligomeric Bri2 BRICHOS (blue), or in the presence of 8% PEG8000 (black) and 50 μM oligomeric Bri2 BRICHOS (orange). The combined chemical shift changes ($\Delta\delta$) determined from the spectra are plotted in the lower panel following the previous color code. Grey circles represent the difference of the chemical shift changes from Bri2 BRICHOS on the dilute phase (blue) and the droplet state (orange), respectively.

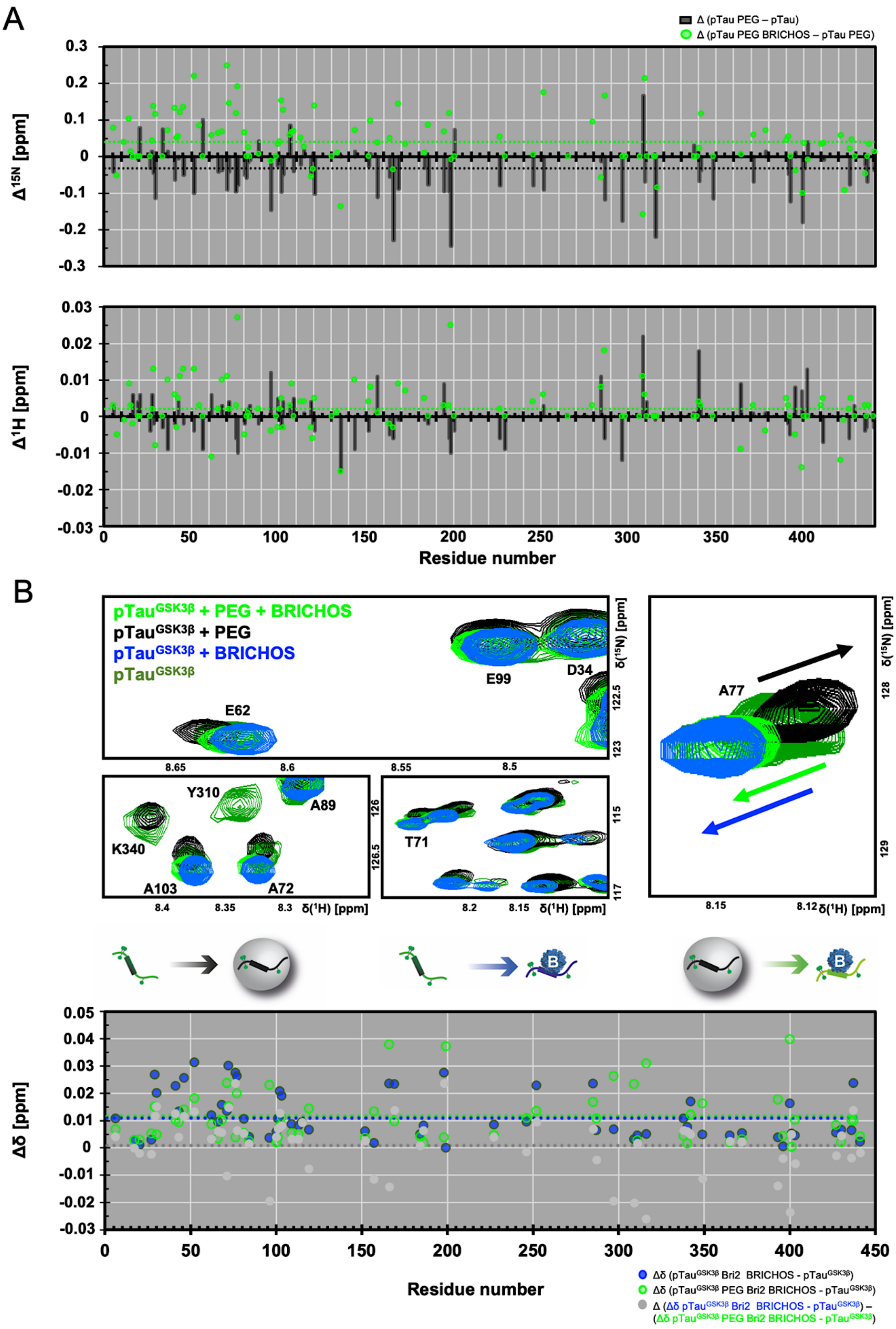


**SI Figure S8. Bri2 BRICHOS counteracts PEG8000-induced chemical shift changes in pTau$^{GSK3\beta}$. (A)** $\Delta^{15}$N and $\Delta^{1}$H chemical shifts of pTau$^{GSK3\beta}$ in the presence of 8% PEG8000 (black bars) or the Bri2 BRICHOS effects on pTau$^{GSK3\beta}$ in PEG8000 (green circles). **(B)** $^{1}$H–

$^{15}$N-HSQC NMR spectra of 100 μM $^{15}$N-labeled monomeric pTau$^{GSK3\beta}$ (red) in the presence of 50 μM oligomeric Bri2 BRICHOS (blue), or in the presence of 8% PEG8000 (black) and 50 μM oligomeric Bri2 BRICHOS (green). The combined chemical shift changes ($\Delta\delta$) determined from the spectra are plotted in the lower panel following the previous color code. Grey circles represent the difference of the chemical shift changes from Bri2 BRICHOS on the dilute phase (blue) and the droplet state (green), respectively.
